# Aberrant Recovery of Timescale-Aligned Amplitude Balance Links to Symptoms and Cognition in Schizophrenia

**DOI:** 10.1101/2025.03.20.644399

**Authors:** Sir-Lord Wiafe, Spencer Kinsey, Najme Soleimani, Raymond O. Nsafoa, Nigar Khasayeva, Amritha Harikumar, Robyn Miller, Vince D. Calhoun

## Abstract

Schizophrenia has long been linked to impaired coordination of brain activity, yet most frameworks overlook two key dimensions: the amplitude of brain signals and the differing timescales on which regions operate. These factors are critical in disorders where neural activity is exaggerated and slowed. In healthy adults, networks compensate for mismatched processing speeds to maintain proportionate activity, but this process is poorly understood in schizophrenia. We developed a timescale-aligned, time-resolved framework that separates temporal distortions from genuine amplitude differences, enabling measurement of amplitude balance between networks across timescales. This approach was applied to large-scale fMRI datasets, including the Human Connectome Project and a multi-site schizophrenia cohort. Patients with schizophrenia showed greater amplitude imbalance, especially during fast fluctuations, along with more frequent re-entry into unbalanced states and slower recovery to stable coordination. We further identified a flexible intermediate state that patients occupied more often, and that predicted better working-memory performance. Across cohorts, amplitude imbalance was associated with greater symptom severity and poorer reasoning ability. These findings provide a new mechanistic view of dyscoordination in schizophrenia grounded in timescale-normalized amplitude dynamics, highlight aberrant recovery of amplitude balance as a core feature of the illness, and suggest that timescale-aligned amplitude imbalance may serve as a promising target for biomarker development.

## 1. INTRODUCTION

Schizophrenia is marked by profound disruptions in large-scale brain coordination, closely tied to symptoms and cognitive deficits^1,2^. Several influential frameworks have been advanced to explain these abnormalities, including theories of dysconnectivity^3,4^, excitation–inhibition imbalance^5,6^, predictive coding failures^7^, and departures from criticality^8^. Each highlights important aspects of network dysfunction, but all largely overlook a fundamental systems challenge: brain networks operate on different internal clocks.

Unimodal regions fluctuate rapidly, transmodal networks integrate more slowly, and neurovascular coupling adds further lags and distortions^9,10^. These temporal mismatches complicate more than just synchrony. They obscure not only synchrony but also amplitude. Two regions engaged in the same operation may appear misaligned in time and thus falsely similar or different in strength. In other words, timescale variability can mask meaningful disparities in network amplitude. This is striking because amplitude is not noise. It often covaries with glucose utilization^11,12^, reflects local synaptic and neuronal activity^13,14^, and shows trait-like stability across individuals^15^. It is therefore functionally relevant to consider not only when regions co-activate, but also the intensity of their activation. We refer to this fluctuation strength simply as network amplitude, which reflects the relative size of BOLD fluctuations after preprocessing.

Despite this relevance, most fMRI studies have prioritized connectivity measures, emphasizing how networks co-vary over time. Approaches such as correlation, sliding-window analysis, phase synchronization, wavelet coherence, and hidden Markov models are intentionally amplitude-invariant, ignoring signal strength^16–19^. Even metrics designed to capture amplitude, such as amplitude of low-frequency fluctuations (ALFF), and their dynamic variants, do not address temporal misalignment across networks^20–22^. As a result, both families of methods underreport amplitude disparities when regions operate on mismatched timescales, leaving a fundamental dimension of brain dynamics hidden in plain sight.

We propose timescale normalization as a systems principle that addresses this challenge. In this view, healthy brain networks compensate for their different intrinsic speeds so that their BOLD amplitudes remain proportionate despite timing mismatches. This compensation should produce relatively homogeneous amplitude disparities and rapid recovery after momentary disruptions^23^. In schizophrenia, however, multiple lines of evidence point to disruptions in processes that depend on timing, such as predictive coding^7^, excitation–inhibition balance^5^, and long-range integration^2^. These abnormalities should weaken timescale normalization, producing more heterogeneous amplitude disparities and slower stabilization after perturbations. Such failures would not only reflect illness expression but also plausibly relate to cognitive decline and symptom burden, because tasks such as working memory, attention, and flexible reasoning rely on rapid coordination across networks^24^. Finally, we hypothesized that abnormalities would be most evident during faster BOLD fluctuations, where timing variability is greatest, and the system is placed under the strongest coordination demands^25^.

To probe this overlooked dimension, we turned to dynamic time warping (DTW), a method that aligns signals by correcting temporal distortions while preserving the size of their fluctuations^26^. In fMRI, DTW has demonstrated greater sensitivity to motor function, robustness to noise, superior test–retest reliability, reduced dependence on global signal regression, and improved detection of group differences (including sex effects and diagnoses such as schizophrenia and autism)^27–29^. We further extended this literature by introducing a warp elasticity metric that quantifies relative BOLD pacing between networks, underscoring DTW’s value in neuroimaging^30,31^. Here, we extend it further, introducing a time-resolved approach that tracks moment-to-moment amplitude disparities between brain networks after correcting for temporal misalignment. These disparities capture the relative proportional balance of activity across networks, a coordination that becomes critical when systems run on different clocks.

We validated the approach through simulations and surrogate analyses, which confirmed its ability to isolate amplitude disparities despite timing variability, and test–retest analyses in a large independent cohort (n = 1,200) established strong reliability. Applied to a multi-site schizophrenia cohort (160 controls, 151 patients), the framework uncovered a mechanistic picture of disrupted network coordination. Static summaries revealed widespread disturbances of timescale-aligned amplitude balance in patients, which were linked to symptom severity and were most evident during faster BOLD fluctuations. Dynamic analyses showed that patients entered unbalanced configurations more often and returned to balance more slowly. We also observed a flexible intermediate mode of coordination that supported cognition: its expression related to stronger working memory and reasoning, while tasks demanding structure, such as verbal learning, benefited from more ordered configurations. Collectively, these findings reveal widespread disturbances of network amplitude balance in schizophrenia, amplified at fast timescales, coupled with slower recovery, and a flexibility–rigidity trade-off that maps onto specific cognitive strengths and weaknesses.

This work introduces a new framework that, for the first time, accounts for the brain’s varying intrinsic timescales when measuring amplitude balance. By separating temporal distortions from true amplitude differences, it provides a nuanced view of large-scale coordination and its breakdown in schizophrenia. Beyond this disorder, the framework offers a generalizable tool for linking brain dynamics to cognition and clinical outcomes, while highlighting biomarkers rooted in the joint principles of temporal and amplitude balance.

## 2. RESULTS

### 2.1. Simulation of DTW sensitivity to non-stationarity and signal bandwidth

We first evaluated DTW as a potential measure of inter-regional amplitude disparity through controlled simulations. These assessed its ability to detect amplitude differences while remaining invariant to phase shifts. Sinusoids with varying amplitude and phase modulations were generated, and DTW was compared to correlation. For DTW, we report both the standard distance and a normalized variant (nDTW), defined as the DTW distance scaled by the alignment-path length, which allows comparability across region pairs and has shown higher sensitivity to group differences^29^. Methods were examined across several DTW 𝛾 parameters and signal bandwidths (Supplementary Material).

DTW successfully aligned original and temporally warped signals, confirming its ability to capture temporal deformations (Fig. 1a). Both DTW and nDTW remained stable under non-stationary phase perturbations, whereas correlation values declined with increasing phase modulation (Fig. 1b). In contrast, when amplitude was perturbed, correlation remained flat, while DTW and nDTW increased with the degree of modulation, indicating selective sensitivity to amplitude disparities (Fig. 1b).

**Fig. 1:**
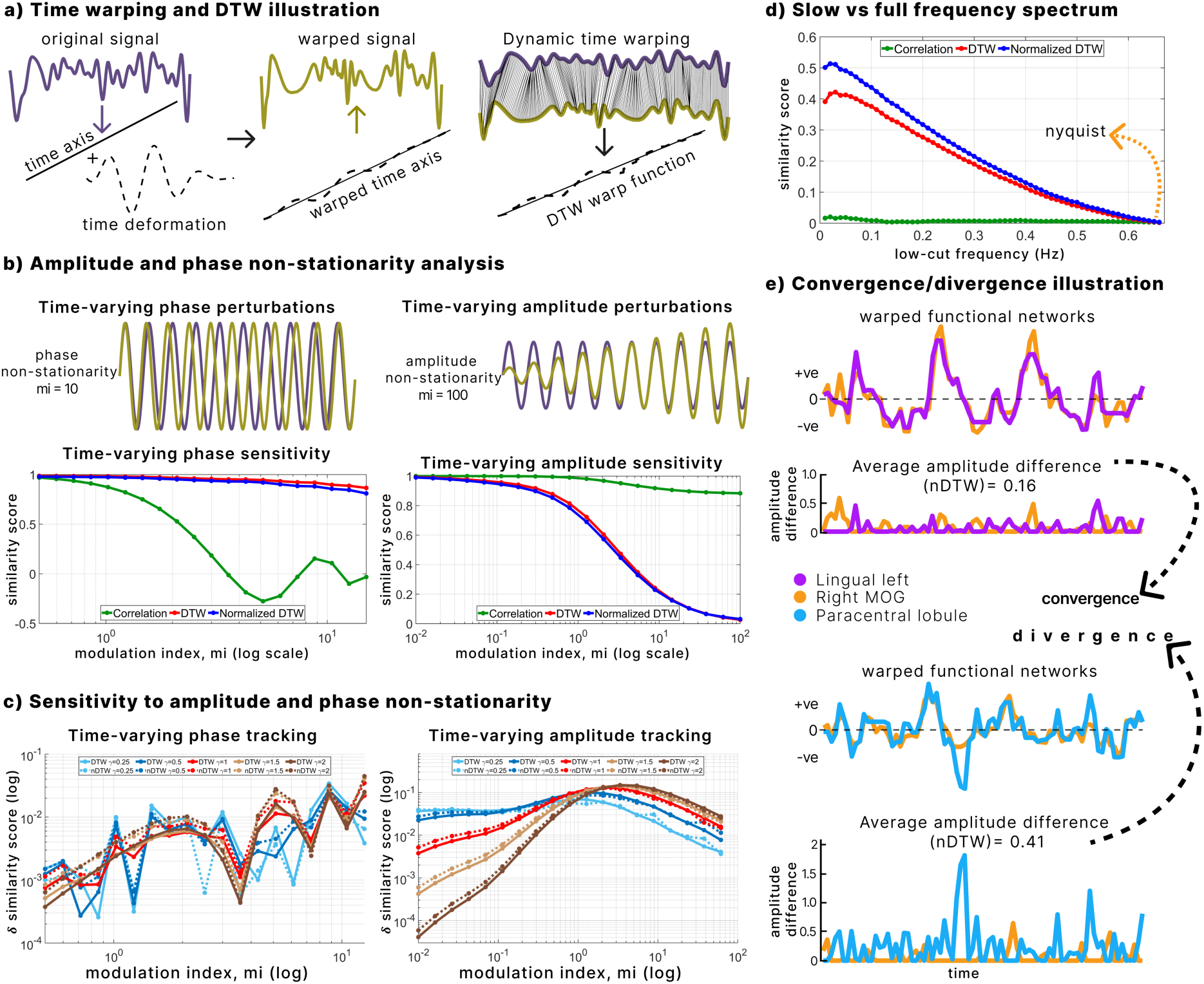
Simulation to analyze non-stationarity and signal bandwidth. **a.** DTW aligns an original signal with its time-warped version, extracting the induced temporal deformation. **b.** Behavior of DTW, nDTW, and correlation under amplitude and phase perturbations. DTW and nDTW detect amplitude disparities but remain largely invariant to phase shifts, unlike correlation. **c.** Sensitivity of DTW and nDTW to the 𝛾 parameter. Smaller 𝛾 values values capture minor amplitude disparities, whereas larger 𝛾 values better capture large disparities. nDTW shows slight phase sensitivity when 𝛾 ≥ 1 (Time-varying phase tracking). **d.** DTW sensitivity across signal bandwidths. Alignment of broadband random signals reveals greater sensitivity to amplitude disparities than correlation, which remains unchanged across bandwidth. **e.** Illustration of convergence and divergence in brain networks. Using nDTW (𝛾 = 1), average amplitude differences are shown for two pairs of networks. Lower nDTW values reflect convergence, while higher values reflect divergence, highlighting imbalances in timescale-normalized amplitudes.

Varying the 𝛾 parameter revealed a trade-off in sensitivity. Smaller 𝛾 values produced stronger responses at low modulation indexes, enhancing detection of subtle amplitude differences, whereas larger 𝛾 values were more responsive at higher indexes, better capturing pronounced disparities (Fig. 1c). Although DTW is generally phase-invariant, nDTW showed slight sensitivity to phase differences at 𝛾 ≥ 1, suggesting it can capture a richer mix of timing and amplitude variation when strict phase invariance is not required.

Bandwidth analyses showed that DTW and nDTW sensitivity increased as signal randomness rose toward the Nyquist frequency, while correlation remained unaffected (Fig. 1d). This indicates that amplitude differences become more detectable in broader frequency ranges.

Finally, Figure 1e illustrates how nDTW (𝛾 = 1) quantifies average aligned amplitude differences. Lower values reflect convergence, where signals remain proportionate, while higher values reflect divergence, where disparities accumulate. This simulation framework establishes DTW, especially nDTW, as a measure capable of distinguishing amplitude-based differences in signals even under temporal distortions, motivating its application to amplitude differences in fMRI data.

### 2.2. Inter-regional timescale-normalized amplitude disparities between schizophrenia and controls across broader fMRI bandwidths using nDTW

The intrinsic brain networks identified using the NeuroMark pipeline were categorized into seven domains: subcortical (SC), auditory (Aud), sensory motor (SM), visual (Vis), cognitive control (CC), default mode (DM), and cerebellum (Cb). To validate nDTW in the context of faster BOLD fluctuations, we extended our analysis to include broader frequency bands (slow-5, slow-4, and slow-3; 0.01–0.198 Hz). We also examined the impact of varying 𝛾 values on test–retest reliability and sensitivity to group differences.

In both frequency ranges (0.01–0.15 Hz and 0.01–0.198 Hz; hereafter F2 and F1 respectively), test–retest reliability improved with higher 𝛾 values, as indicated by lower mean ranks (Fig. 2a). 𝛾 values of ≥1.5 demonstrated significantly higher reliability compared to correlation (Wilcoxon signed-rank test, FDR corrected p < 0.05). Because physiological noise correction in the fMRI data reduces within-subject variability^32^, the improved reliability at higher 𝛾 values likely reflects reduced sensitivity to physiological noise.

**Fig. 2:**
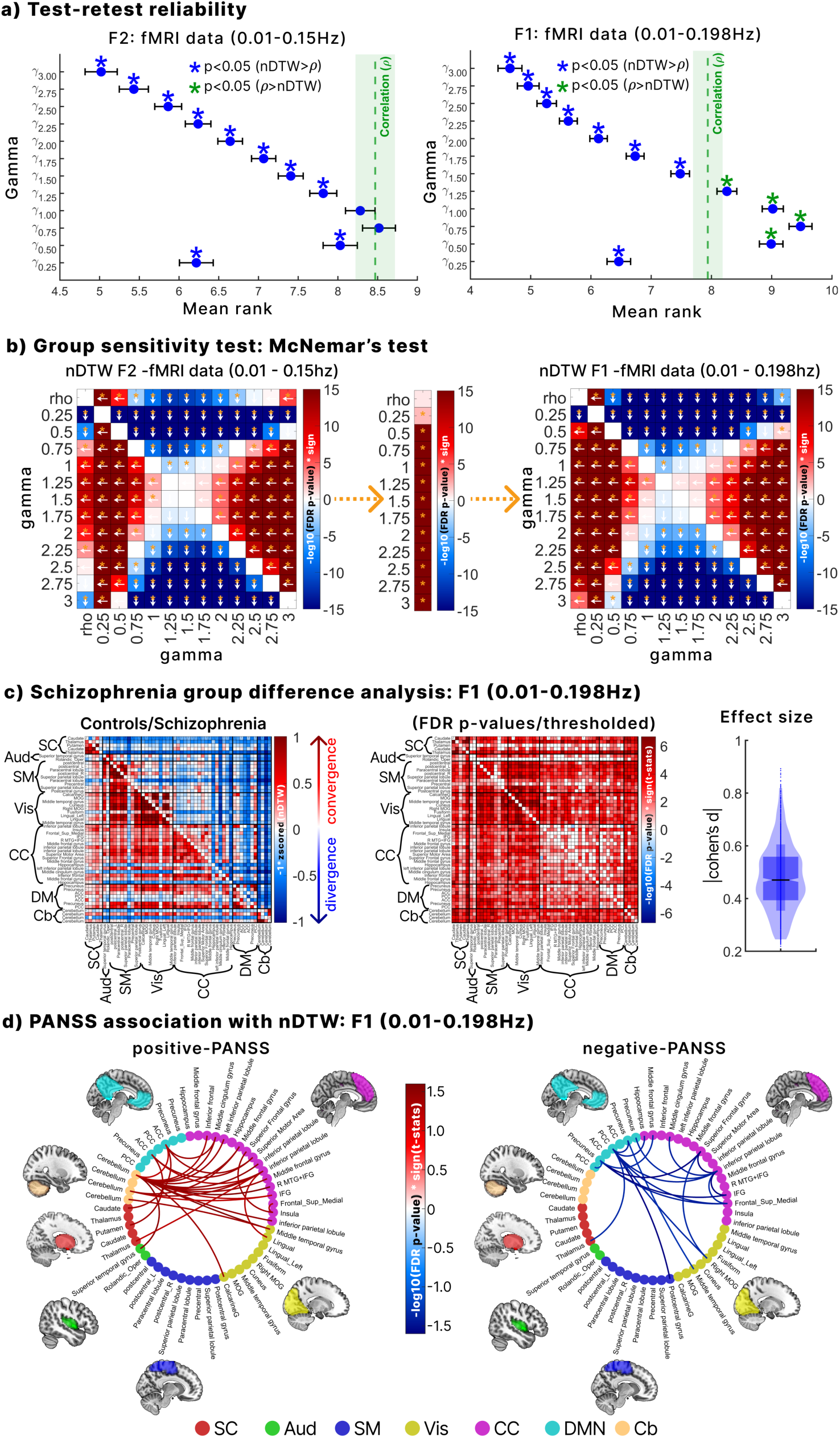
Reliability and group difference sensitivity. **a**. Test–retest reliability of nDTW across 𝛾 values in the HCP dataset, with correlation as a benchmark. Lower mean rank indicates higher reliability. Values of 𝛾 ≥ 1.5 and 𝛾 = 0.25 showed significantly greater reliability than correlation (Wilcoxon signed-rank test, FDR-corrected p < 0.05) in both F1 and F2 frequency configurations. **b.** Sensitivity of 𝛾 values for distinguishing schizophrenia from controls across frequency ranges. McNemar tests compared all γ values in F1 and F2, and between F1 and F2. Arrows indicate the more sensitive parameter; yellow asterisks mark significant results (FDR-corrected p < 0.05). 𝛾 = 1.25 and 𝛾 = 1.5 provided the strongest group discrimination. Sensitivity improved from F1 to F2 for all 𝛾 values, but not for correlation. **c.** Group differences at 𝛾 = 1.5. nDTW values were z-scored, inverted (so convergence is positive, divergence negative), and averaged across groups. Controls showed stronger convergence across many network pairs, while patients showed divergence. Significant group effects (FDR < 0.05, GLM controlling for sex, age, site, and motion) are marked, with effect sizes (Cohen’s d > 0.2) supporting practical relevance. **d.** The association of nDTW with symptoms. Significant links between network pairs and both positive and negative PANSS scores (GLM, Poisson distribution, FDR-corrected p < 0.05) are shown, adjusted for sex, age, site, motion, and CPZ (patient-only GLMs). These associations highlight relationships between amplitude disparities and symptom severity in schizophrenia.

Figure 2b shows that 𝛾 values of 1.5 and 1.25 achieved the greatest sensitivity for detecting group differences across both frequency bands, outperforming other 𝛾 values and correlation. However, since 𝛾 = 1.25 has significantly lower test–retest reliability than the correlation benchmark in F1 range (Fig. 2a), 𝛾 = 1.5 was selected as the optimal parameter for further analysis. Additionally, nDTW consistently detected more group differences in the broader F1 frequency range than in F2 across all 𝛾 values, whereas correlation did not. This demonstrates that nDTW is particularly sensitive to clinically relevant group differences during faster BOLD fluctuations while maintaining high reliability at 𝛾 = 1.5.

Figure 2c illustrates that controls (lower triangle) showed lower amplitude disparity (greater convergence), whereas individuals with schizophrenia (upper triangle) exhibited higher disparity (greater divergence) across multiple network pairs. Group comparisons confirmed significantly more convergence in controls across several networks (FDR-corrected, Fig. 2c). Effect sizes were substantial (Cohen’s d range 0.19–0.89, median = 0.49), highlighting the practical significance of these disparities.

Finally, significant associations between amplitude disparity and PANSS scores were identified. Higher positive PANSS scores were linked to more balanced/homogenous disparity between cerebellar networks and the DM, CC, and superior temporal gyrus (Aud), as well as with Vis and SC networks. Higher negative PANSS scores were mainly associated with greater disparity between CC, DM, and Vis networks.

### 2.3. Effects of including faster BOLD fluctuations on amplitude disparity

We evaluated the effect of expanding the frequency range from F2 to F1 using nDTW, with paired-sample t-tests conducted separately in controls and patients. Normality of nDTW scores was confirmed using Shapiro–Wilk tests, with values close to 1 across network pairs (Fig. 3a). After FDR correction, controls exhibited significantly lower amplitude disparity (greater convergence) in F1 compared to F2, whereas schizophrenia showed higher disparity (greater divergence) in F1 than in F2 across several network pairs. These findings suggest that including faster BOLD fluctuations enhances balance in controls but amplifies imbalance in schizophrenia, thereby increasing sensitivity to group differences.

**Fig. 3:**
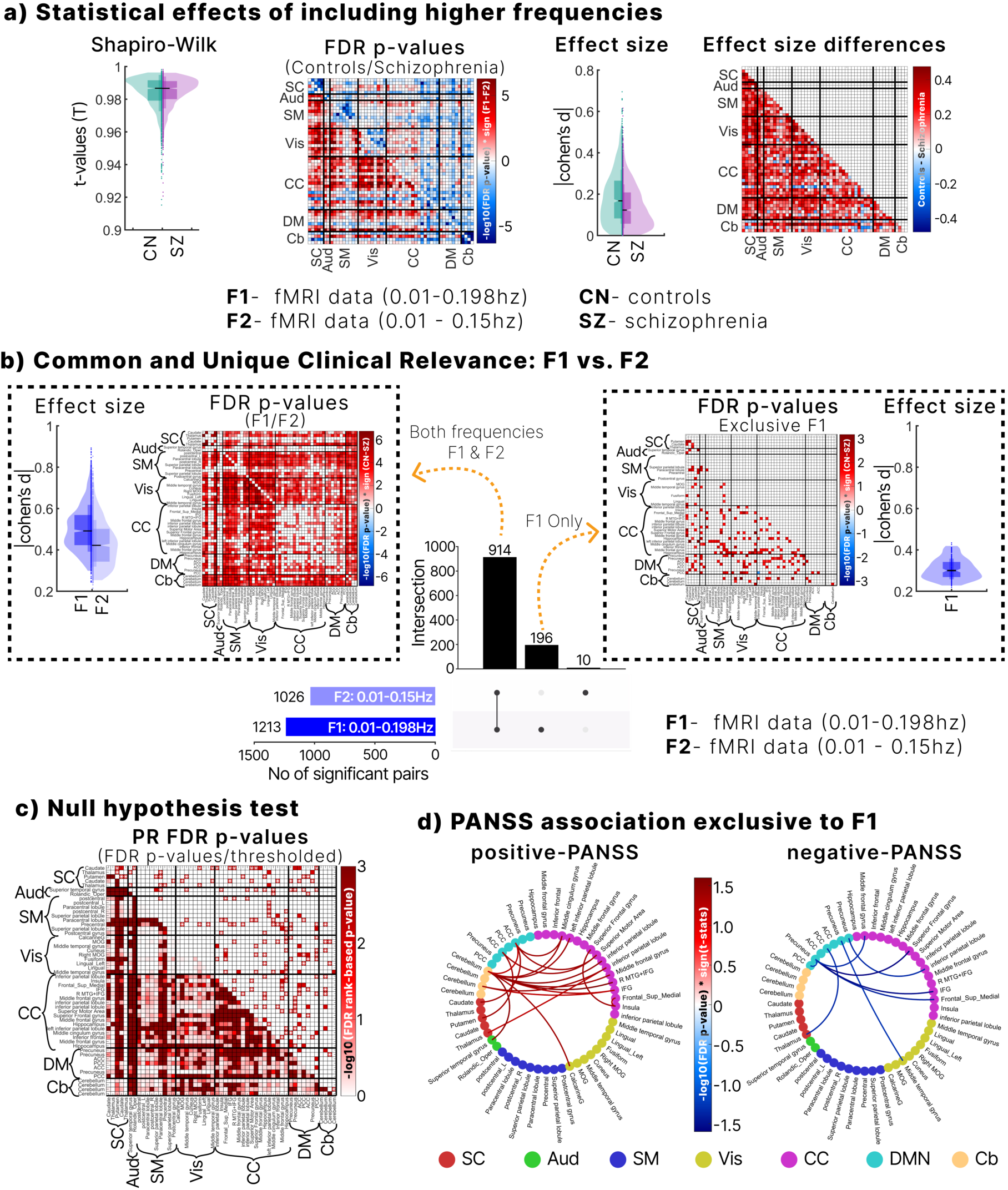
Clinical significance of high frequency fMRI data inclusion. **a**. Timescale-normalized amplitude imbalances across frequency ranges. Normality was confirmed with Shapiro–Wilk tests (T ≈ 1), validating paired t-tests. Controls showed greater convergence, whereas schizophrenia patients showed divergence when higher frequencies were included (F1 vs F2). Cohen’s d (∼0.4) indicated enhanced separability between groups. **b.** Group differences across network pairs from a generalized linear model controlling for sex, age, motion, and site. In total, 914 pairs were significant in both bands, with F1 showing larger effect sizes and stronger relevance. An additional 196 pairs were uniquely significant in F1 (effect sizes > 0.2), compared with only 10 unique pairs in F2. **c.** Null hypothesis testing using phase randomization confirmed that the significant brain network pairs uniquely identified in F1 are statistically relevant. **d.** Associations between normalized nDTW and PANSS scores were exclusively observed in the F1 frequency range, with no significant associations found in F2.

Effect sizes were modest overall (median Cohen’s d = 0.18 for controls and 0.15 for schizophrenia). However, faster frequencies amplified distinctions between groups, as indicated by larger effect size differences across multiple network pairs (Fig. 3a). Upset plot analysis identified 914 significant pairs across both frequency ranges, with 196 unique to F1 and only 10 unique to F2, underscoring the added sensitivity of F1 (Fig. 3b).

Among pairs significant in both ranges, F1 (lower triangle) showed higher practical relevance than F2 (upper triangle), with unique F1 pairs displaying Cohen’s d > 0.2. To further validate F1-specific effects, we generated 1,000 surrogate datasets through phase randomization within the 0.15–0.198 Hz range. Several pairs remained significant (Fig. 3c), confirming that faster fluctuation effects are statistically robust and reflect non-stationary amplitude processes rather than noise.

Finally, symptom analyses revealed that associations with PANSS were clearer in F1 than in F2. Positive PANSS correlated with disparities linking the cerebellum to SC, DM, CC, Aud, and Vis networks, while negative PANSS was primarily associated with CC–DM connections. These associations were absent in F2, suggesting that faster BOLD fluctuations improve sensitivity to clinically relevant symptom links.

### 2.4. Dynamics of time-resolved DTW

To identify recurring states of amplitude imbalance dynamics, we applied k-means clustering with three clusters on the time-resolved DTW, including both controls and patients. An elbow plot (Fig. 4c) confirmed that three clusters were optimal, determined by the derivative of the within-sum of squared distances. For visualization, we subtracted the median of all clusters and multiplied the distribution by −1 so that positive values represent convergence and negative values indicate divergence (Fig. 4a). The clusters were classified as follows: state 1, a convergent state; state 3, a divergent state; and state 2, a mixed state showing neither strong convergence nor divergence.

**Fig. 4:**
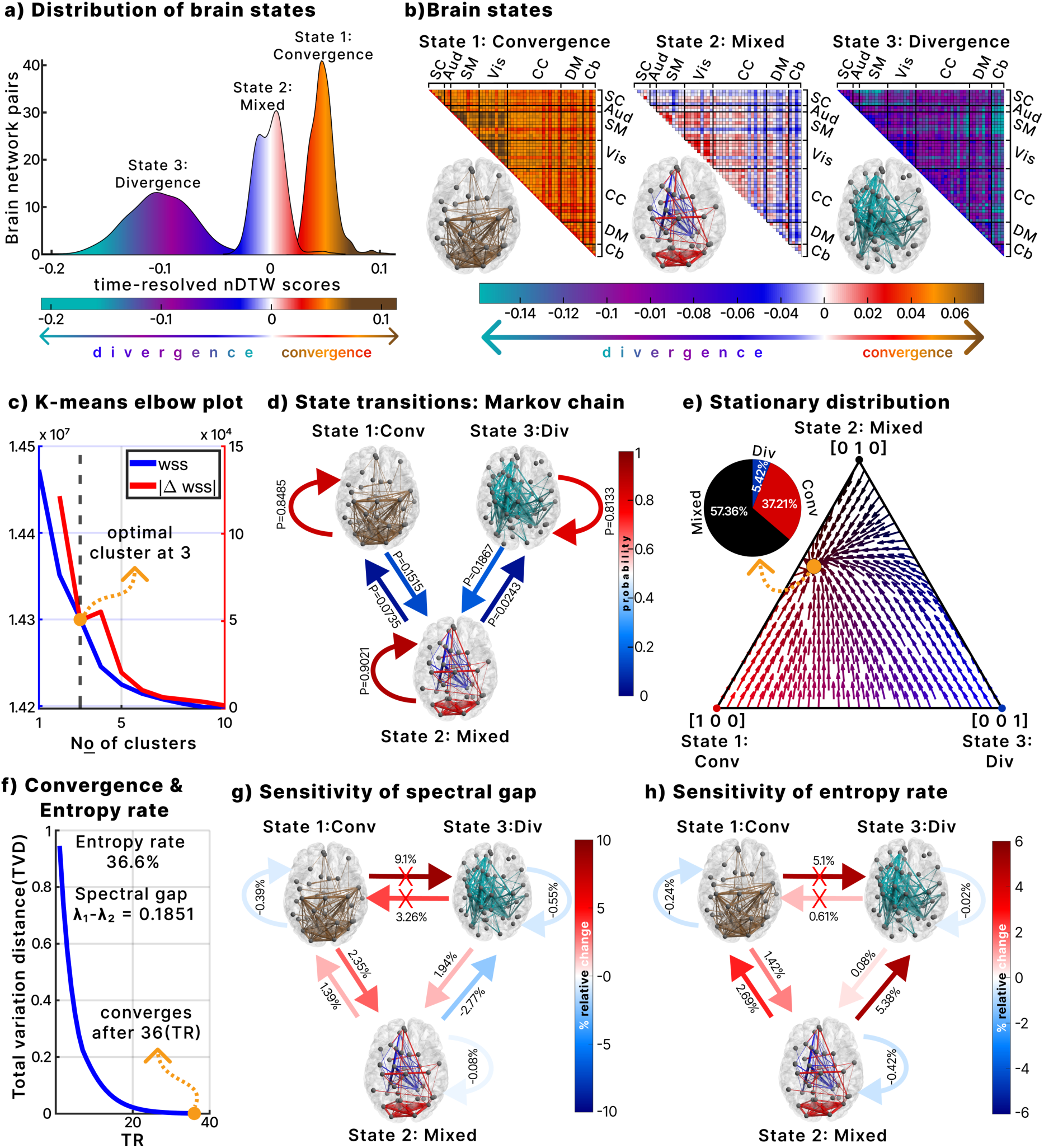
Dynamics of time-resolved DTW. **a.** Distribution of recurring states identified with k-means clustering on time-resolved DTW. Centroids were normalized so positive values represent convergence and negative values represent divergence. Three states emerged: convergent, divergent, and mixed. **b.** Brain state matrices with axial thumbnails. Convergent and divergent states show the top 5% of strongest connections, while the mixed state shows the top 2.5% most convergent and 2.5% most divergent connections. **c.** Elbow plot of within-sum of squares (WSS). The first inflection point in the derivative of WSS indicated three optimal clusters. **d.** State transition probabilities between the three states. **e.** Stationary distribution of the Markov chain, showing that long-run dynamics are shaped primarily by the mixed and convergent states. **f.** Mixing time of the dynamics, with convergence rate further characterized by the spectral gap and entropy rate. **g.** Perturbation of state transitions and effects on the spectral gap, showing changes in convergence rate. **h.** Perturbation of state transitions and effects on the entropy rate.

Figure 4b shows the network representations for each state. In convergent state 1, the most convergent pairs (top 5 percent) predominantly involved sensory networks including Vis, SM, and Aud. Divergent state 3 featured the most divergent pairs (top 5 percent), mainly between the Cb and other networks. Mixed state 2 combined both dynamics, with the top 2.5 percent most convergent and the top 2.5 percent most divergent pairs.

A Markov chain was constructed to model state transitions (Fig. 4d). Mixed state 2 had the highest self-retention probability (0.9021), followed by convergent state 1 (0.8485) and divergent state 3 (0.8133). No direct transitions occurred between divergent and convergent states, consistent with a regime in which shifts proceed through the intermediate mixed state.

The stationary distribution of the Markov chain (Fig. 4e) showed that mixed state 2 accounted for 57.36 percent of dynamics, convergent state 1 for 37.21 percent, and divergent state 3 for 5.42 percent. This highlights the predominance of mixed and convergent states in maintaining amplitude balance.

We further analyzed these dynamics by calculating the entropy rate and the rate of return to the stationary distribution (spectral gap). The entropy rate was 36.6 percent, where 100 percent represents complete randomness and 0 percent full order. This indicates a balance between randomness and order, with a modest bias toward order (Fig. 4f). The spectral gap was 0.1851, corresponding to a mixing time of 36 timesteps (72 seconds, TR = 2). The mixing time was derived from the farthest distribution point in the Markov chain’s state space, specifically the state [0, 0, 1] (completely divergent), as shown in Fig. 4e.

Finally, we explored how entropy and convergence rate were influenced by transition probabilities using an exploratory perturbation analysis (Fig. 4g–h). Increasing self-retention in any state reduced both the convergence rate and entropy. Adding transitions between the mixed state and other states (excluding mixed to divergent) enhanced both rates. By contrast, direct transitions between divergent and convergent states were not considered because none were observed in the data (Fig. 4d). Increasing transition probability from mixed to divergent states extended trajectories away from the stationary distribution, slowing return to stability. Although this increased entropy, the results show that higher entropy does not universally accelerate recovery; its effect depends on the specific dynamic transitions within the state space.

### 2.5. Group differences between controls and schizophrenia

We compared groups on metrics derived from time-resolved DTW, including mean dwell time, occupancy rate, state transition probabilities, stationary distribution, spectral gap, mixing time, and entropy rate. Patients with schizophrenia were more often associated with the mixed (state 2) and divergent (state 3) configurations, whereas controls predominantly engaged with the convergent state (state 1). Specifically, patients showed longer mean dwell times in the mixed and divergent states, while controls spent more time in the convergent state (Fig. 5a). Occupancy rates showed a similar pattern, with higher mixed and divergent occupancy in patients and greater convergent occupancy in controls (Fig. 5b).

**Fig. 5:**
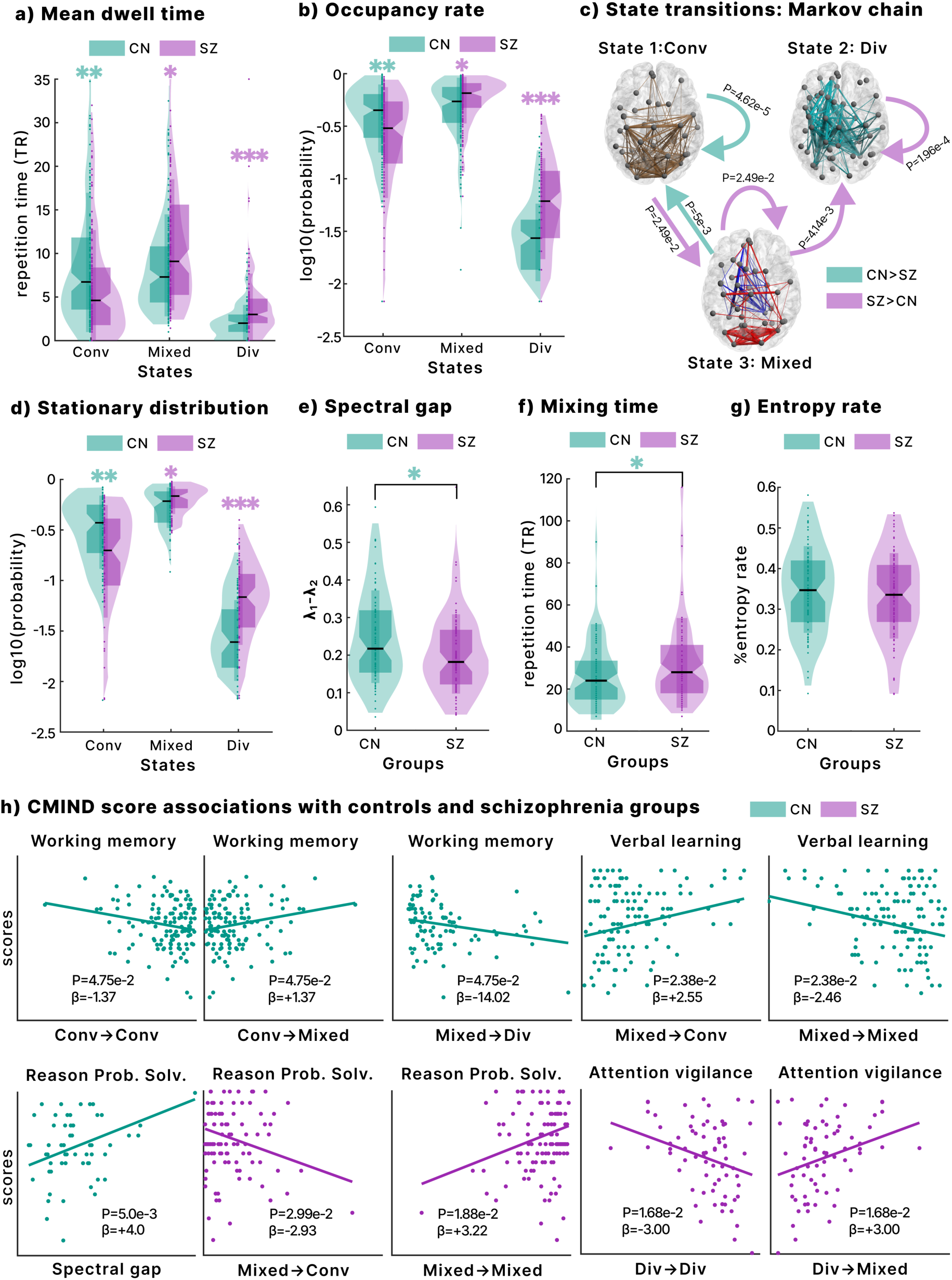
Group differences and cognitive score association. **a.** Mean dwell time in each state, showing longer time in mixed and divergent states for schizophrenia and in the convergent state for controls**. b.** Occupancy rate, with the same group pattern as dwell time. **c.** State transition probabilities, highlighting higher rates from convergent to mixed and mixed to divergent states in schizophrenia. **d.** Group differences in the stationary distribution of the Markov chain. **e.** Spectral gap, reflecting convergence rate, was higher in controls, indicating faster stabilization. **f.** Mixing time, representing the steps to reach the stationary distribution, was shorter in controls than in schizophrenia. **g**. Entropy rate, capturing order–flexibility balance, showed no group differences. **h.** Associations between cognitive scores and nDTW metrics. In controls, working memory and verbal learning are related to state transitions, and reasoning to the spectral gap. In schizophrenia, reasoning, problem-solving, and attention vigilance were linked to different transitions and state retention. *All analyses used generalized linear models controlling for sex, age, site, motion, and CPZ (for patient-only GLMs). Results were FDR-corrected (significance threshold = 0.05).* 𝑝 < 0.05 *(*),* 𝑝 < 0.01 *(**),* 𝑝 < 0.001 *(***)*.

The stationary distribution confirmed that schizophrenia dynamics were more strongly shaped by the mixed and divergent states, while controls were more strongly shaped by the convergent state (Fig. 5d). Transition probabilities further highlighted group differences: controls were more likely to move from mixed to convergent states, whereas patients more often transitioned from convergent to mixed, from mixed to divergent, and showed higher self-retention in both mixed and divergent states (Fig. 5c).

Controls also exhibited a higher spectral gap, reflecting a faster rate of return to the stationary distribution (Fig. 5e). Starting from the fully divergent configuration ([0, 0, 1]) and using a tolerance of 10^-3^, controls had a shorter mixing time, suggesting a quicker recovery to a stable amplitude-balanced configuration. By contrast, entropy rates did not differ significantly between groups, indicating that the faster recovery observed in controls is likely explained by transition structure rather than overall system randomness.

### 2.6. Associations with cognition in controls and schizophrenia

We tested associations between time-resolved DTW metrics and Computerized Multiphasic Interactive Neurocognitive System (CMINDS) scores using generalized linear models (GLM) for controls and patients separately (Fig. 5h).

In controls, working memory was positively associated with transitions from convergent to mixed states and negatively with convergent self-retention and mixed-to-divergent transitions, suggesting that entering the mixed state supports memory performance. Verbal learning was positively associated with transitions from mixed to convergent states and negatively with convergent-to-mixed transitions, indicating that the convergent state facilitates verbal learning. Reasoning and problem-solving were positively associated with the spectral gap, linking faster convergence rates with higher cognitive ability.

In schizophrenia, reasoning and problem-solving showed the opposite pattern: they were negatively associated with transitions from mixed to convergent states but positively with self-retention in the mixed state. Attention vigilance was positively associated with transitions from divergent to mixed states and negatively with self-retention in the divergent state. These findings indicate that dynamic transitions contribute differently to cognition in patients, where entry into the mixed state supports vigilance, while persistent divergence impairs attention.

## 3. DISCUSSION

This study introduces a new framework to assess timescale-normalized amplitude imbalances between brain networks using fMRI. By focusing on the amplitude of BOLD signals, referred to here simply as network amplitude, and applying a time-resolved DTW approach, we provide a timing-invariant readout of amplitude differences that accounts for the brain’s varying temporal scales. Most prior fMRI research has emphasized synchrony and connectivity, leaving amplitude underexplored despite its physiological and functional relevance. Our framework addresses this gap by isolating genuine amplitude disparities from temporal distortions, yielding a clearer view of how large-scale systems coordinate and how that coordination fails in schizophrenia. This study provides the first mechanistic account of schizophrenia that jointly corrects for mismatched timescales and amplitude disparities, revealing a fundamental principle of brain coordination.

Using the recently developed nDTW metric^29^, we extended analysis beyond the conventional 0.01–0.15 Hz band to include faster BOLD fluctuations. Time-resolved DTW revealed three intuitive brain states: convergence (lower disparity), divergence (higher disparity), and a mixed state representing flexible balance. Simulations confirmed the robustness of nDTW in capturing aligned amplitude differences while remaining insensitive to timing and phase shifts, and empirical application showed that state dynamics of amplitude disparity were meaningfully associated with both clinical symptoms and cognition.

Static analyses revealed that controls displayed stronger convergence, whereas patients with schizophrenia exhibited greater divergence, indicating a tendency toward imbalance across networks. These results suggest that healthy brains maintain more proportionate amplitude balance globally, while schizophrenia is marked by greater heterogeneity and imbalance. This aligns with prior reports of altered physiological and metabolic processes in schizophrenia^33,34^.

A key observation was that including faster BOLD fluctuations enhanced sensitivity to group differences. In controls, these higher frequencies were associated with more proportionate amplitude balance, while in schizophrenia, they were linked to greater imbalance. Moreover, significant differences between groups emerged exclusively when faster fluctuations were included, and a phase-randomized null model confirmed these effects were not attributable to noise. Although high-frequency BOLD signals have often been dismissed as artifacts^35^, other work has identified them as meaningful sources of neural variability^36–38^. Our findings provide a novel perspective by showing that faster fluctuations improve the detection of schizophrenia-related imbalances, likely by capturing rapid non-stationary processes overlooked in conventional analyses.

We also observed strong associations with clinical symptoms. Positive symptoms correlated with altered cerebellar interactions, particularly with auditory, DM, CC, SC, and visual networks. The cerebellum is a recognized predictive hub^39^, and central to the “cognitive dysmetria” theory, which attributes schizophrenia to impaired coordination among prefrontal, thalamic, and cerebellar regions^40^. Our results support and extend this theory by showing that cerebellar-related amplitude disparities track positive symptom severity. Interestingly, lower divergence correlated with more severe positive symptoms, suggesting maladaptive or compensatory cerebellar involvement.^41^

Negative symptoms were instead linked to disparities within higher-order processing networks such as DM and CC. These findings are consistent with evidence that negative symptoms, characterized by reduced motivation, pleasure, and emotional expression, are linked to high-level cognitive processing^42^. Prior studies have connected impaired physiology and metabolism to negative symptoms like depression^43,44^, and a recent study found that improvements in metabolic measures, achieved through prefrontal cortex stimulation, were associated with reductions in negative symptoms^45^. Our results complement this literature by providing a timing-invariant, amplitude-based marker that relates amplitude imbalance to symptom severity. Notably, many of these symptom associations depended on the inclusion of faster BOLD fluctuations, underscoring the importance of extending analyses beyond traditional frequency ranges.

Dynamic analyses revealed three recurring states. The convergent state represented lower amplitude disparity, the divergent state higher disparity, and the mixed state intermediate dynamics. Transitions occurred only between mixed and either convergent or divergent states, never directly between convergence and divergence, suggesting that the mixed state serves as an adaptive intermediary. The stationary distribution showed the mixed state dominated, accounting for 57% of system dynamics, with convergence at 37% and divergence at 5%.

These results resonate with brain criticality theory, which posits that the brain operates near a critical point between order and disorder^46^. The prevalence of the mixed state, combined with a moderate entropy rate of 36.6%, supports the idea that brain dynamics balance stability and flexibility, with a slight bias toward order but remaining within the “zone of criticality”^47^. Indeed, critical entropy serves as an indicator of brain criticality^48^, a state that helps the brain maintain flexibility and adaptability in its functions. We interpret these links cautiously as consistency rather than a direct test of criticality.

A key dynamic metric in our study is the spectral gap, which measures how quickly the system returns to its stationary distribution. Perturbation analysis showed that increasing self-retention reduced both return rate and entropy, since greater predictability comes at the expense of flexibility central to criticality theory. In general, higher return rates were associated with increased entropy, except for transitions from the mixed to divergent state, underscoring the role of specific transition pathways in shaping stability. State-space analysis revealed that trajectories from the divergent state required the longest path to reach stability, explaining why an excess of mixed-to-divergent transitions slowed recovery. Controls exhibited significantly higher spectral gaps and shorter mixing times than patients, indicating faster stabilization despite similar entropy levels. These results suggest that schizophrenia disrupts transition pathways supporting efficient recovery, and they highlight that high entropy alone does not guarantee effective information integration or cognitive performance^49^.

Cognitive flexibility was highlighted by associations with the mixed state. In controls, working memory improved with transitions from convergence into the mixed state and declined with both self-retention in convergence and shifts from mixed to divergence, indicating that the mixed state, reflecting flexible amplitude balance, supports higher-order functions such as working memory^50^. In schizophrenia, reasoning and problem-solving were negatively related to mixed-to-convergent transitions but positively linked to sustained retention in the mixed state and to divergence-to-mixed transitions. Vigilance also declined with self-retention in divergence yet improved with divergence-to-mixed transitions, underscoring the importance of flexible exits from imbalanced configurations. These results suggest that disrupted transitions into and out of the mixed state may underlie cognitive impairments in schizophrenia.

However, the mixed state may not be optimal for all tasks. In controls, mixed-to-convergent transitions were positively associated with verbal learning, whereas extended retention in the mixed state was negatively related. Given that verbal learning benefits from structured training^51^, consistent with the ordered nature of the convergent state, it appears that while the mixed state promotes flexibility, tasks requiring structured cognition rely more on stability. Overall, these findings underscore the need to balance flexibility and stability for optimal cognitive performance.

Group-level analysis revealed that controls predominantly engaged in the convergent state, while schizophrenia subjects showed greater involvement in the mixed and divergent states. Metrics such as mean dwell time, fraction rate, state transitions and stationary distribution confirmed these patterns. The increased presence of divergent states in schizophrenia aligns with the disorder’s characteristic disruption in neural dynamics^52^. Furthermore, as transitions toward the mixed state were found to decrease the rate of return to stability, the excessive engagement of the mixed state in schizophrenia may reflect an overly sustained flexibility that potentially undermines cognitive processes such as working memory.

Although this framework provides a robust approach for quantifying timescale-normalized amplitude imbalances, several considerations should be noted. First, BOLD amplitude is inherently a relative measure influenced by preprocessing choices, ICA scaling, and spatial heterogeneity in signal intensity; therefore, our metric reflects aligned differences in signal strength rather than absolute physiological quantities, and caution is warranted when linking these disparities to underlying biology. Second, although our sample sizes (HCP: 827; fBIRN: 311) support stable estimation of dynamic metrics, studies with smaller cohorts may place greater emphasis on the static amplitude-disparity measure, which is less sensitive to sample size than transition-based quantities such as spectral gap or entropy rate. Despite these considerations, the framework has clear translational promise: identifying maladaptive amplitude-imbalance trajectories may inform patient-specific markers of state instability, guide future neuromodulation targets, and provide a scalable analytic tool for other psychiatric and neurological conditions.

In summary, this framework demonstrates that schizophrenia is characterized by widespread inter-regional amplitude disparities, most pronounced during rapid neural processing. Positive symptoms were linked to cerebellar-related imbalances, while negative symptoms were associated with higher-order network disruptions. Dynamic analyses revealed flexible but inefficient reliance on mixed states, slower recovery to stability, and links between state transitions and cognition. These findings provide a new and nuanced mechanistic view of schizophrenia, situating its disruptions in the joint dimensions of temporal alignment and amplitude balance. Beyond schizophrenia, this framework establishes a generalizable approach for linking brain dynamics to cognition and clinical outcomes, and it highlights potential biomarker targets grounded in fundamental principles of temporal and amplitude balance. Importantly, the framework also points toward translational opportunities: in the Supplementary Material, we outline a potential therapeutic application using photobiomodulation guided by directional DTW. By targeting maladaptive trajectories of amplitude imbalance, such approaches may offer novel avenues for intervention in psychiatric illness.

## 4. METHODS

### 4.1. Dynamic time warping as a measure of amplitude disparity

Dynamic time warping is a widely used signal processing algorithm that aligns signals with nonlinear temporal deformations by applying adaptive “elastic” transformations to achieve a meaningful similarity measure^26,53^. When temporal deformations complicate phase or timing relationships, it becomes logical to isolate these aspects and concentrate solely on the amplitude of the signals. DTW tackles this by resolving temporal dissimilarities, optimally aligning signal pairs to minimize a distance metric and bring them to a shared temporal scale, leaving amplitude as the primary source of disparity.

The algorithm computes a distance cost matrix that captures the cumulative cost of aligning pairwise elements of the sequences and identifies the optimal warping function to adjust their time axes, minimizing a distance metric. To enhance computational efficiency and ensure meaningful alignments, particularly when guided by domain knowledge, a window size constraint is applied to limit the maximum allowable warping between indices in the time series^26^.

For discrete signals 𝑥 and 𝑦 of lengths 𝑁 and 𝑀 respectively, the distance cost matrix, 𝐶𝑀*_DTW(i,j)_*, is initialized as follows:

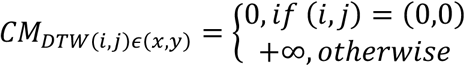

The cells in 𝐶𝑀*_DTW(i,j)_* that are within the allowable window, 𝑤, can be defined as:

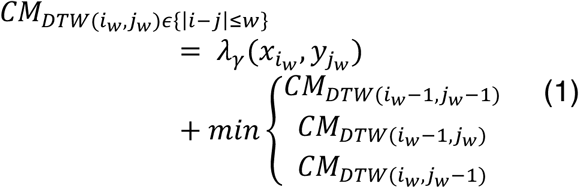

Following the parameterization of the DTW algorithm recently proposed^54^, let 𝛾𝜖ℝ^+^ denote that 𝛾 is a positive real number. Then, the generalized distance metric is given as:

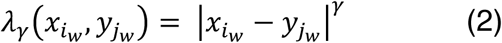

Where 𝛾 > 1 emphasizes large amplitude disparity, 𝛾 < 1 highlights small amplitude disparity and 𝛾 = 1 provides a balance between both^54^.

The optimal warping path or function 𝜑, with length 𝐿, derived from DTW, is the path that belongs to the set of all possible warping functions Π, where the cumulative distance metric between corresponding points in signals 𝑥 and 𝑦 is minimized. It can be expressed as:

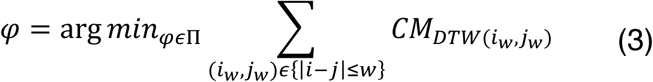

Where 𝐿 ≥ max (𝑀, 𝑁)

The distance cost 𝐷 in DTW, which measures the phase/timing-invariant amplitude disparity between the signal pair, is computed as the sum of distance metric along the warping path. This cost is expressed as:

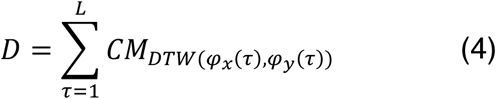

Where 𝜏 is the time index for the warp function, 𝜑_𝑥_(𝜏) is the warping path along signal 𝑥, and 𝜑_𝑦_(𝜏) is the warping path along signal 𝑦.

Notably, when the distance metric with 𝛾 = 2 is chosen the DTW distance cost becomes:

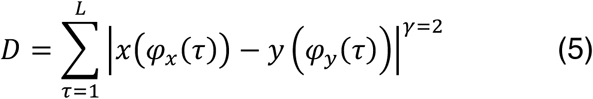

This represents the conventional energy (sum of squared aligned differences) of the difference signal 𝑥(𝜑_𝑥_(𝜏)) − 𝑦 (𝜑_𝑦_(𝜏)), and the normalized DTW distance cost 𝐷*_n_* represents the power (mean squared aligned difference) of the difference signal, as given below:

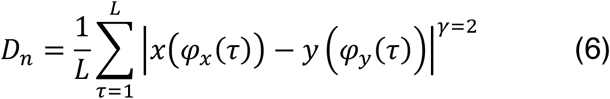

This positions the algorithm as a measure of as a measure of timing-invariant amplitude disparity between two signals. Leveraging the 𝛾 −parameterization, DTW and nDTW tune sensitivity to amplitude differences 𝛾 > 1 upweights large disparities, 𝛾 < 1 upweights small disparities and 𝛾 = 1 offers the perfect balance.

We define convergence as relatively low nDTW (smaller time-invariant amplitude difference), and divergence as high nDTW (larger time-invariant amplitude difference).

### 4.2. Allowable window size for fMRI application

The two main challenges of DTW are its computational cost, with a time complexity of 𝑂(𝑀 × 𝑁), and the issue of singularities^26^, where a single time point in one signal is unnaturally matched to a large portion of the other signal. Both challenges are mitigated by constraining the alignment to a window size that limits the maximum window of allowable alignment^55^. Given the focus on temporal deformations such as leads/lags and nonlinear dynamics like stretching and shrinking, selecting an appropriate window size to capture relevant temporal deformations in fMRI data is crucial. In our previous study using the proposed warp elasticity approach, we demonstrated that stretching and shrinking dynamics could introduce anti-correlations^31^. Considering that anti-correlations may also arise from lead/lag relationships, it is intuitive to choose a window size that spans the length of the signal required to produce maximum anti-correlations (−1). This approach has also been validated as optimal in prior fMRI studies^28^. The longest wavelength yielding an anti-correlation of −1 corresponds to the lowest frequency component (0.01𝐻𝑧), which equates to a 100-second window. Achieving an anti-correlation of −1, requires a time shift of half the wavelength (50 seconds). For the DTW algorithm to capture relevant temporal deformations in fMRI data, two 50-second windows are sufficient to account for deformations in both directions, resulting in a total window length of 100 seconds^28^, corresponding to the low-cutoff frequency of 0.01𝐻𝑧. However, since the filter design for the data, commonly Butterworth, is typically nonideal, we instead recommend selecting a window size based on the spectral characteristics of the activity signal, determined by the −3𝑑𝑏 point of the low-frequency cutoff of the fMRI signal^18,30,31,56^. The equation for determining the window size is:

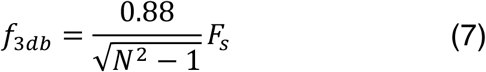

Where 𝑁 represents the number of points in the window, 𝐹_s_ denotes the sampling frequency, and 𝑓_3*db*_ is the low cutoff frequency.

### 4.3. Time-resolved DTW

To derive a time-resolved evolution of amplitude imbalances, we modify the nDTW measure by eliminating its summation and normalization components. This yields a time-resolved amplitude mismatch metric, 𝐷*_tr_*(𝜏), for the aligned signals, ensuring that its average corresponds to the original nDTW value. The formulations are as follows:

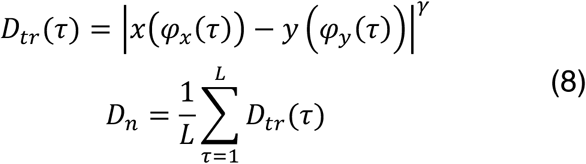

However, the DTW algorithm results in aligned signal lengths 𝐿 that vary across different signal pairs, complicating comparisons between brain network pairs. To ensure that the length of the time-resolved metric equals the signal length and averages to the nDTW, it is necessary to interpolate 𝐷*_tr_*(𝜏) with the warped time scale (𝜏) to have the same length as the original time index (𝑡), 𝑁 or 𝑀 (noting that 𝑁 = 𝑀). This interpolation must preserve the integrity of the metric, particularly the first derivatives of the warping functions, which are crucial for capturing the varying temporal scales of fMRI time series as established in our previous study^31^.

To address this, we employed the piecewise cubic Hermite interpolating polynomial (PCHIP) method for interpolating 𝐷*_tr_*(𝜏) to the original signal length 𝑁. PCHIP was selected for its ability to preserve the shape and monotonicity^57^ of the original warping functions, thereby maintaining the temporal dynamics of the fMRI signals. Unlike standard cubic splines, PCHIP avoids introducing artificial oscillations and overshoots^57^, ensuring the fidelity of the amplitude mismatch measurements. Additionally, PCHIP guarantees continuous first derivatives^57^, aligning with our requirement to maintain the derivatives of the warping functions. This interpolation ensures that the interpolated metric 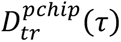 accurately averages to the 𝐷*_n_*, facilitating consistent and comparable analyses across all brain network pairs.

The interpolated time-resolved DTW metric is thus defined as:

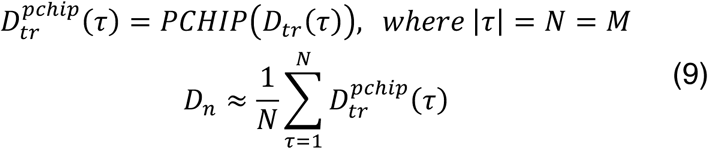

To validate our approach, we generated 10,000 independent and identically distributed (i.i.d.) Gaussian signal pairs, each sampled from a standard normal distribution 𝒩(0,1). Calculating the percentage relative difference between the nDTW and the average time-resolved DTW yielded a median percentage difference of 0.20% (Supplementary Material).

This methodology ensures that the time-resolved metric retains its temporal dynamics and derivative continuity while standardizing its length. Consequently, preserving 𝐷*_n_* within the time-resolved measure maintains its reliability, thereby enhancing the robustness of our analysis.

### 4.4. Resting-state fMRI data

The first dataset used is from the Function Biomedical Informatics Research Network (fBIRN) study including resting fMRI data collected from schizophrenia patients and controls^58^. Scans were collected at a repetition time (TR) of 2 seconds. The study led to a total of 160 controls with an average age of 37.04 ± 10.86 years, ranging from 19 to 59 years. Among these, 45 were female and 115 were male. Additionally, there were 151 patients diagnosed with schizophrenia, with an average age of 38.77 ± 11.63 years, ranging from 18 to 62 years. In this group, 36 were female and 115 were male. The typical controls and schizophrenic patients were meticulously matched in terms of age, gender distribution, and mean framewise displacement during scans (age: p = 0.18; gender: p = 0.39; mean framewise displacement: p = 0.97). Schizophrenic patients were in a clinically stable condition during the time of their scans. The fBIRN dataset is used in all the clinical analysis performed in this study.

The second dataset used in our study is the resting-state fMRI dataset collected from 827 subjects via the Human Connectome Project (HCP) database^59,60^. We utilized both session scans acquired using a Siemens Skyra 3T scanner with a multiband accelerated, gradient-echo echo-planar imaging (EPI) sequence. The scanning parameters were set to a TR of 0.72s, 72 slices, an echo time (TE) of 58ms, and a flip angle of 90°. A voxel size of 2 × 2 × 2 mm was used to acquire 1200 time points of fMRI data for each subject. The HCP dataset was used for our test-retest analysis.

### 4.5. Resting-state fMRI pre-processing

fMRI data requires extensive preprocessing to correct for various sources of noise and artifacts before analysis. The preprocessing steps commonly applied in fMRI studies include slice timing correction, realignment, spatial normalization, and spatial smoothing^61^. Following typical preprocessing^62^, we implemented the NeuroMark pipeline, a fully automated spatially constrained ICA on the preprocessed fMRI data^63^. Using the neuromark_fMRI_1.0 template (available in GIFT at http://trendscenter.org/software/gift or http://trendscenter.org/data), we generated 53 intrinsic connectivity networks (ICNs) for each subject. These ICNs are grouped into brain functional domains, including subcortical, auditory, sensorimotor, visual, cognitive control, default mode, and cerebellum.

### 4.6. Resting-state fMRI post-processing

To enhance the quality of these ICNs, we implemented detrending and despiking methods to eliminate drifts, sudden fluctuations, and significant artifacts that may have persisted after the initial preprocessing stages. The time series of the ICNs were bandpass filtered within two frequency ranges: F1 (0.01-0.198Hz) and F2(0.01-0.15Hz). An infinite impulse response filter was designed using the butter function in MATLAB and applied via the *filtfilt* function to ensure zero phase shifts and preserve phase information. The optimal filter order was estimated using the *Buttord* function in MATLAB, which returns the lowest order of the Butterworth filter with no more than 3dB ripple in the passband and at least 30dB of attenuation in the stopband. Finally, we performed z-scoring on the ICNs. The MATLAB version used was MATLAB R2022a for all steps of the analysis.

### 4.7. Test-retest reliability

Typically, test-retest reliability is evaluated using the intra-class correlation coefficient (ICC). However, the ICC relies on a linear parametric ANOVA model that assumes separability and additivity^64^. This makes ICC unsuitable for DTW applications, as DTW generates a distance metric based on nonlinear warping, where pairwise distances are interdependent^28,29^ and thus lack decomposable variance components. We utilize the HCP dataset, which includes two sessions and a large sample size of 827 subjects, facilitating robust test-retest analysis.

#### 4.7.1. Within-subject variability

We adopt the approach of a previous study by employing a straightforward method to assess within-subject variability: a paired sample t-test between sessions for each brain network pair^28^. The absolute t-value is reported as a measure of how much the differences between the two sessions deviate from zero, where lower absolute t-values indicate higher within-subject consistency. The formula for calculating the absolute t-value between two sessions of a single network pair across all subjects is provided below:

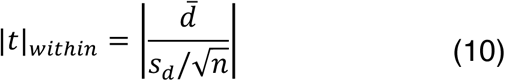

Where 𝑑̅ is the mean difference between two sessions, 𝑠*_d_* is the standard deviation of the differences, and 𝑛 is the total number of subjects.

#### 4.7.2. Between-subject variability

To assess between-subject variability, we developed a non-parametric equivalent of the absolute t-value. First, we calculate the differences between the means of each subject’s distribution to capture variability across the group. Since the direction of these differences is not of primary interest, we take their absolute values, resulting in a distribution skewed toward zero. This transformation allows us to perform a non-parametric test to evaluate how a non-parametric central tendency measure, such as the median, deviates from zero. Unlike the parametric approach, which scales the central tendency measure by the standard deviation, our non-parametric method scales it by the interquartile range to account for the skewed distribution. The formula for the non-parametric absolute variability measure for a single session is as follows:

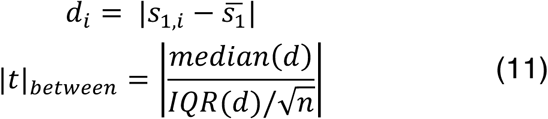

Where 𝑑*_i_* is the absolute difference between subject 𝑖’s measure in session 1 (𝑠_1,*i*_) and the group mean for session 1 (𝑠͞_1_), 𝐼𝑄𝑅(𝑑) is the interquartile range of the differences 𝑑, and 𝑛 is the total number of subjects.

This non-parametric approach provides a robust measure of between-subject variability that is less influenced by outliers and non-normal distributions, thereby enhancing the reliability of our variability assessments.

#### 4.7.3. Composite test-retest reliability score

To further summarize these two approaches to give a wholistic view of test-retest reliability, we compute a composite score of test-retest as follows:

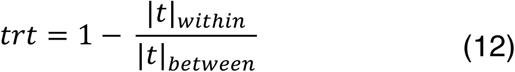

Scores approaching 1 indicate higher test-retest reliability.

In our study, we compared 𝑡𝑟𝑡 scores across various gamma values of nDTW against correlation as a benchmark. To achieve this, we ranked the 𝑡𝑟𝑡 scores and performed Wilcoxon signed-rank tests between each gamma value’s 𝑡𝑟𝑡 scores and the correlation benchmark across all brain network pairs, following a critical difference computation approach.

### 4.8. Group difference and clinical association analyses

We employed generalized linear models for both group difference analyses and associations with clinical scores. All GLM analyses controlled for sex, age, site, and mean frame displacement. For clinical association analyses conducted within the schizophrenia group, chlorpromazine-equivalent (CPZ) antipsychotic dose was included as a covariate. P-values were adjusted using the false discovery rate method, with significance set at a threshold of 0.05. For group difference analyses and computerized multiphasic interactive neurocognitive system scores, a normal distribution was assumed, whereas a Poisson distribution was applied for PANSS score associations due to the positive integer and skewed nature of the fBIRN dataset’s PANSS scores. Effect sizes were calculated using Cohen’s d.

### 4.9. Group difference sensitivity test

To evaluate the sensitivity of various gamma values of nDTW and correlation in distinguishing between controls and individuals with schizophrenia, we conducted McNemar’s sensitivity tests comparing each gamma value and correlation against each other. Following the GLM group difference analyses, we counted the number of significant brain network pairs (FDR corrected p < 0.05) for each gamma value (nDTW) and for correlation. We then compared the unique and shared significant pairs between each gamma and correlation using McNemar’s test, with p-values adjusted via FDR correction across all comparisons.

These sensitivity tests were performed separately for each frequency range, F1 and F2. Additionally, we compared F1 and F2 across all gamma values and correlation metric to assess how transitioning from the lower frequency range F2 to the broader F1 range affects group difference sensitivity.

### 4.10. Validation of high fMRI frequency results

High-frequency fMRI signals are vulnerable to various sources of noise and artifacts, including physiological factors like respiratory and cardiac activity, as well as non-neuronal influences such as head motion^65^. Comprehensive preprocessing steps followed by our use of ICA help mitigate these concerns. ICA effectively separates independent components, allowing us to identify and remove artifact-related components from the data. Additionally, we included head motion parameters as covariates in our GLM analyses to ensure that detected group differences in high-frequency signals are not attributable to head motion effects.

We conducted several additional analyses to further validate our findings within the F1 frequency range.

#### 4.10.1. Statistical effects on groups with the inclusion of fast BOLD fluctuations

We aimed to evaluate the impact of transitioning from the F2 to the broader F1 frequency range on both control and schizophrenia groups. Since F1 and F2 comprise the same dataset differentiated only by frequency range, a paired sample t-test was appropriate for this analysis. To validate the use of the paired t-test, we first assessed the normality of nDTW scores for each group between F1 and F2 using the Shapiro-Wilk test, ensuring that t-values were close to 1.

Subsequently, we conducted paired t-tests separately for the control and schizophrenia groups to examine the effects of transitioning from F2 to F1, with p-values adjusted using the FDR correction. Effect sizes for these comparisons were calculated using Cohen’s d. Additionally, to highlight the differential impact of transitioning from F2 to F1 between groups, we computed the differences in Cohen’s d effect sizes between controls and schizophrenia. This comparison aimed to determine whether the transition to the broader F1 frequency range enhances or diminishes group-specific distinctions.

Furthermore, we identified statistical group differences between schizophrenia and controls that were present in F1 but not in F2 by analyzing the shared and unique significant brain network pairs from our GLM analysis (see Section 3.7). We also highlighted their corresponding effect sizes, demonstrating that significant group differences are more pronounced in the F1 frequency range.

#### 4.10.2. Null hypothesis test

The phase randomization (PR) technique generates surrogate fMRI data by preserving the correlation between brain network pairs^66^. This method is suitable for fMRI null hypothesis testing as it assesses non-stationarity by capturing information beyond linear and static correlations^66^. In our previous study^29^, we employed PR to evaluate the statistical relevance of both DTW and nDTW in fMRI data, demonstrating that both metrics rejected the null hypothesis for several brain region pairs. This validation confirms that the DTW and nDTW methods are sensitive to capturing information beyond traditional correlation measures.

In this study, we utilize PR to assess the statistical relevance of nDTW in capturing non-stationary amplitude changes specifically due to the inclusion of higher frequency data (0.15–0.198 Hz). The following steps were followed to generate 1,000 surrogate fMRI datasets:

I Phase Randomization: Apply PR to the F1 frequency range data (0.01–0.198 Hz) to create surrogate data, denoted as 𝑍.
II Filtering 𝑍: Use the designed filter applied to achieve original F2 fMRI data (see section 3.4.) on 𝑍 to extract the 0.01–0.15 Hz frequency range of the surrogate data, resulting in surrogate data 𝑌.
III Isolating Higher Frequencies: Subtract 𝑌 from 𝑍 to obtain the surrogate data residual high frequency 0.15–0.198 Hz, denoted as 𝑋.
IV Full surrogate data: Add 𝑋 to the original F2 fMRI data (0.01–0.15 Hz) to generate surrogate data with phase randomization affecting only the residual 0.15–0.198 Hz frequency range.

This process generates surrogate datasets where randomization impacts solely the residual 0.15–0.198 Hz frequency range. A high-pass filter was not applied on 𝑍 to obtain the 0.15–0.198 Hz surrogate data because non-ideal filters will artificially create frequency multiples at the edges (∼0.15 Hz).

We performed a Wilcoxon rank-sum test (FDR-corrected p < 0.05) to compare nDTW scores from the original fMRI data with those from surrogate data. This test assessed whether the significant brain network pairs distinguishing controls from schizophrenia, identified exclusively in the F1 range, also rejected the null hypothesis.

#### 4.10.3. PANSS association exclusive to F1

We identified significant brain network pairs associated with symptom scores that were exclusive to the F1 frequency range and not present in the F2 range.

### 4.11. Amplitude imbalance dynamics

#### 4.11.1. Brain state estimation

In time-resolved fMRI studies, k-means clustering is frequently utilized to identify recurring brain states within the temporal dynamics of the data^67^. The DTW algorithm inherently anchors the first and last time points during alignment, potentially introducing bias by disproportionately influencing brain states at the temporal extremes. To mitigate this bias, we excluded the first and last 5 time points from the time-resolved DTW metrics prior to clustering. This exclusion reduces alignment-induced artifacts, enabling the clustering algorithm to more accurately capture intrinsic transitions between brain states.

We concatenated the time-resolved DTW metrics from all subjects in the fBIRN dataset into a single feature matrix, where each time point from every subject serves as an individual sample and each brain network pair constitutes a distinct feature. We implemented the k-means algorithm with cluster numbers ranging from 1 to 10, setting the maximum number of iterations to 10,000 and using 20 random initializations to ensure robust convergence. The city-block (Manhattan) distance metric was selected for its demonstrated robustness in handling high-dimensional data^68^. To determine the optimal number of clusters, we generated an elbow plot based on the within-cluster sum of squares (WSS). The optimal cluster number was identified as 3, corresponding to the first significant inflection point in the WSS curve’s first derivative.

#### 4.11.2. Markov chain & stationary distribution

We quantified the transitions between brain states by counting the number of times each subject moved from one state to another and normalizing these counts across each state to construct individual transition probability matrices. This represents the probability of a subject transitioning from one state to another and further provides insights into the Markov chain of state dynamics. To gain a comprehensive understanding of the general transition patterns independent of specific groups, such as patients and controls, we aggregate the probability transitions across all subjects in the fBIRN dataset.

Furthermore, the stationary distribution of the Markov chain represents the long-term behavior of the system, indicating the proportion of time the system spends in each state once it has converged to equilibrium^69^. This distribution serves as an indication of a “stable point” when analyzing systems from the perspective of their Markov chains rather than traditional state-space representations. For the stationary distribution to be unique and ensure convergence from any initial state, the Markov chain must be ergodic^70^. A Markov chain is considered ergodic if it is irreducible (it is possible to get from any state to any other state, aperiodic (the system does not cycle in a fixed pattern), and positive recurrent (the expected return time to each state is finite).

The stationary distribution 𝜋 is computed by solving the following system of linear equations:

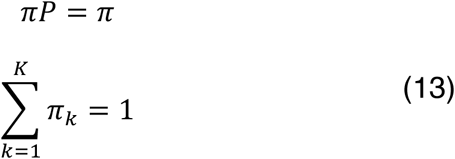

Where 𝜋 = [𝜋_1_, 𝜋_2_, …, 𝜋*_k_*] is the stationary distribution vector, 𝑃 is the transition probability matrix of the Markov chain, 𝐾 is the number of distinct states from K-means (3 in our study).

#### 4.11.3. Convergence analysis

How fast a chain takes to reach the stationary distribution gives a similar equivalent meaning to how fast a system takes to reach its stable point from any given state of the system. The spectral gap gives the rate of this convergence to equilibrium^69^. A larger spectral gap implies faster convergence rate to the stationary distribution. It is computed as follows:

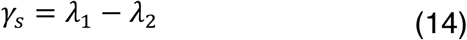

Where 𝜆_1_, and 𝜆_2_ are the largest and second largest eigen values of the Markov chain respectively. When the chain is ergodic, 𝜆_1_ = 1.

Beyond the spectral gap, the mixing time provides a more comprehensive measure of how quickly the chain approaches equilibrium^69^. The mixing time quantifies the number of steps required for the chain’s state distribution to approach the stationary distribution within a specified tolerance level. A prevalent method for determining the mixing time leverages the total variation (TV) distance, which offers a rigorous metric for assessing the divergence between two probability distributions. The total variation distance at time step 𝑡 is defined as:

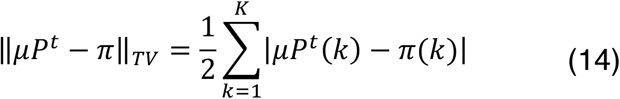

Where 𝜇 is the initial probability distribution, 𝑃*^t^* represents its 𝑡-th power of transition matrix, indicating the state probabilities after 𝑡 transitions, 𝜋(𝑘) is the stationary probability of state 𝑘 and 𝐾 is the total number of distinct states.

The mixing time, denoted as 𝑡*_MT_*, is the smallest positive integer 𝑡 for which the TV distance falls below a predefined tolerance level 𝑡𝑜𝑙:

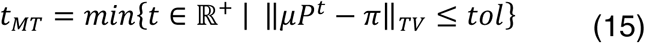

For our analysis, we set the tolerance level to 𝑡𝑜𝑙 = 10^-3^. We also find the mixing time using the initial probability distribution 𝜇 farthest from the stationary distribution.

These metrics, spectral gap and mixing time, are instrumental in understanding the dynamic behavior of brain network states. The spectral gap provides insight into the inherent convergence rate of the system, while the mixing time offers a tangible measure of how quickly the system reaches equilibrium. Together, they facilitate a comprehensive analysis of amplitude imbalances and their temporal evolution within the fMRI data, enhancing the robustness and reliability of our findings.

#### 4.11.4. Entropy rate

We investigated how the trDTW metric informs the flexibility or rigidity of the brain’s dynamic amplitude imbalance configuration by analyzing the entropy rate. The entropy rate is defined as the limit of the average entropy per step as the number of steps approaches infinity. For an ergodic Markov chain, the entropy rate in bits can be mathematically expressed as:

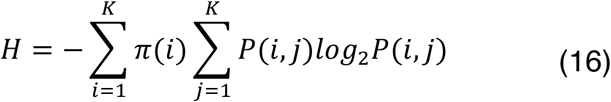

Where 𝜋(𝑖) is the stationary probability of being in state 𝑖 and 𝑃(𝑖, 𝑗) is the transition probability from state 𝑖 to state 𝑗.

The maximum entropy rate for 𝐾 = 3 states is:

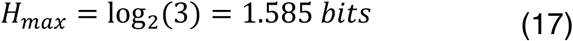

In our study, we present entropy rate as a percentage of 𝐻*_max_* to enhance interpretability.

The entropy rate provides insight into the balance between order and randomness within the amplitude imbalance dynamics. A low percentage entropy rate indicates strong order in the dynamics. Conversely, a high percentage entropy rate signifies a high degree of randomness, suggesting unpredictable amplitude dynamics between brain networks.

#### 4.11.5. Impact of transition probabilities on Markov chain convergence and entropy rate

To evaluate how specific transitions within the Markov chain influence the convergence and entropy rates of the dynamics, we systematically perturbed the aggregated transition matrix. This process involved incrementally increasing the probability of individual transition cells and subsequently renormalizing the corresponding rows to ensure that each row’s probabilities sum to one. Consequently, the targeted transition’s probability increased while the probabilities of other transitions within the same row decreased.

For each perturbed transition, we recalculated both the spectral gap and the entropy rate to determine their percentage relative changes compared to the original values. This perturbation procedure was applied to every transition cell across 10,000 random samples, with perturbation magnitudes ranging from 10^-1^to 10^-3^ on a logarithmic scale to ensure uniform sampling. To maintain comparability between transitions, each sample employed a single perturbation magnitude uniformly across all transition cells.

### 4.12. Time-resolved DTW group difference: schizophrenia vs controls

To assess the clinical relevance of the time-resolved DTW measure, we conducted group difference analyses on several derived metrics: state transitions, stationary distribution, spectral gap, mixing time, and entropy rate (see Section 3.11). Unlike the aggregated approach in Section 3.11, these metrics were calculated individually for each subject. To ensure the uniqueness of these results, we excluded subjects with non-ergodic state transitions.

Additionally, we included mean dwell time and occupancy rate in our group comparisons. Mean dwell time quantifies the average duration a subject remains in a particular state once entered^71^, reflecting the persistence of states within the Markov chain. Occupancy rate represents the percentage of time a subject spends in each state^71^, indicating the relative prevalence of states during the observation period.

Using these subject-specific metrics, we distinguished schizophrenia patients from control subjects and performed group difference analyses employing a GLM, as described in Section 3.8. FDR correction is applied to each metric across all three identified states.

### 4.13. CMINDS score association

Similarly, we employed a GLM, as detailed in Section 3.8, to assess associations between each metric and CMINDS scores—including speed of processing, attention vigilance, working memory, verbal learning, visual learning, and reasoning/problem solving—within the fBIRN dataset. To account for potential group effects, separate GLM analyses were conducted for each group. For transition probabilities, we applied FDR correction to control for multiple comparisons across all transitions.

## 5. ACKNOWLEDGMENT

**Sir-Lord Wiafe:** Conceptualization, Formal analysis, Methodology, Visualization, Writing –original draft. **Spencer Kinsey:** Methodology, Validation, Writing – review & editing. **Najme Soleimani:** Validation, Writing – review & editing. **Raymond O. Nsafoa:** Validation, Supervision. **Nigar Khasayeva:** Validation, Writing – review & editing. **Amritha Harikumar:** Validation, Writing – review & editing. **Robyn Miller:** Validation, Methodology, Supervision, Writing – review & editing. **Vince Calhoun:** Funding acquisition, Validation, Methodology, Resources, Supervision, Writing – review & editing.

## 8. DATA & CODE AVAILABILITY STATEMENT

The codes for all our analyses in MATLAB language can be accessed through GitHub (https://github.com/Sirlord-Sen/time_resolved_dynamic_time_warping). The data was not collected by us and was provided in a deidentified manner. The IRB will not allow sharing of data or individual derivatives as a data reuse agreement was not signed by the subjects during the original acquisition.

