## supplementary material for "Aberrant Recovery of Timescale-Aligned Amplitude Balance Links to Symptoms and Cognition in Schizophrenia" (from another version of this preprint: version 2, 2025-09-22)

***Supplementary simulations***

I. *DTW illustration*

This simulation demonstrates the application of non-linear time warping and the ability of dynamic time warping (DTW) to recover the applied temporal distortion. With a fixed sampling interval of $T_{r}=1$ second, and a signal of length $L=200$, the base signal is created by applying bandpass filtering to white noise (i.i.d Gaussian, $\mathcal{N}(0,1)$) within the frequency range $\left[ 0.01 0.1 \right] Hz$. After filtering, the signal is standardized using z-score normalization.

Simultaneously, a warping function is generated from bandpass-filtered noise in a narrower frequency band $\left[ 0.01 0.02 \right] Hz$. This function is scaled using a maximum stretch value of 15, yielding a warp factor of 15. The resulting warping function, $w(t)$, is then added to the original time vector $t$ to create a non-linearly modulated time axis:

$$t_{warp}=t+w(t)$$

To simulate realistic temporal distortions, the base signal is interpolated along the warped time vector $t_{warp}$, resulting in the warped signal. This warped signal is also normalized using z-score normalization to maintain consistency. Finally, DTW is applied to the pair of signals to compute an optimal alignment between the original and warped signals. This process yields an estimated warping function, $\hat{w}(t)$, which is then compared to the true warping function $w(t)$. Through this comparison, the simulation demonstrates the capability of DTW in identifying and recovering non-linear time distortions.

II. *Amplitude and phase non-stationarity*

These simulations evaluate how different modulations—phase and amplitude—affect the similarity between a base sine wave and its perturbed versions, using DTW alongside correlation measures.

In the phase modulation analysis, a base sine wave is defined by:

$$s\left( t \right)=sin(2\pi ft)$$

with a frequency of $f=0.01Hz$, generated over 1500 samples using a sampling interval of $T_{r}=0.72$ seconds. Phase perturbations are introduced by adding a slow sine modulation of frequency $f_{m}=0.0001 Hz$, so that the modulated signal becomes

$$s_{ph}(t)=sin\left( 2\pi ft+a sin\left( 2\pi f_{m}t \right) \right)$$

where the modulation scale $a$ varies logarithmically from 0.5 to 15. After both the original and phase-modulated signals are standardized using z-score normalization, similarity is quantified by computing a correlation similarity index and a DTW-based similarity index. The DTW similarity index is defined as

$$score_{similarity}=1-\frac{D}{D_{upper}}$$

where $D$ is the DTW distance between the z-scored signals, and $D_{upper}$ is an upper-bound DTW distance computed by applying an extremely large phase perturbation (i.e. scrambling the phase by adding a term like $1000000 sin\left( 2\pi f_{m}t \right)$). Additionally, DTW is computed for five different gamma values, $\gamma\in\{0.25, 0.5, 1, 1.5, 2\}$, which weight the local distance measure in the DTW algorithm.

The amplitude modulation analysis follows a similar structure. Here, the base sine wave is modulated in amplitude to yield a signal defined by

$$s_{amp}(t)=s\left( t \right)\cdot\left( 1+ a sin\left( 2\pi f_{m}t \right) \right)$$

with the modulation scale $a$ varying logarithmically from 0.01 to 100. After applying z-score normalization, the same similarity measures—correlation, and both DTW-based similarity indices—are evaluated. An upper-bound DTW distance is established by generating a maximally perturbed signal through extreme amplitude scaling, and subsequent DTW distances are normalized against this worst-case reference. This normalization ensures that comparisons remain consistent across the different gamma values.

III. *Frequency broadband sensitivity*

These simulations assess the impact of low pass filtering on white noise signals by comparing similarity measures computed using DTW and Pearson correlation. In this setup, two independent white noise signals, denoted as $x(t)$ and $y(t)$, are generated at a sampling interval of $T_{r}=0.72$ seconds over 1000 time points. After detrending to remove any linear components, each signal is passed through a low-pass Butterworth filter whose cutoff frequency $f_{c}$ is varied from $0.01 Hz$ to $0.66 Hz$ —approaching the Nyquist frequency.

For the DTW-based similarity, two types of distances are computed: a raw DTW distance $D$ and a normalized DTW distance obtained by dividing $D$ by the length of the warping path. To facilitate meaningful normalization, an upper-bound DTW distance is first established. This is done by filtering the signals using an extreme cutoff of 0.67 Hz over 1000 realizations, which provides a worst-case reference $D_{upper}$ (and its normalized counterpart $D_{upper}^{norm}$). For any filtered signal pair, the DTW similarity indices are then defined as:

$$score_{DTW}=1-\frac{D}{D_{upper}} and score_{DTW_{norm}}=1-\frac{D/L}{D_{upper}^{norm}}$$

ensuring that a similarity value of 1 corresponds to perfect alignment.

In parallel, the Pearson correlation is calculated between the z-scored filtered signals, providing an alternative measure of similarity.

Finally, for each cutoff frequency and across 100 realizations, these measures are averaged. This robust averaging allows for an insightful evaluation of how varying broadband of signals influences the similarity between stochastic signals, revealing trends in the DTW and correlation metrics as the filter cutoff varies.

***Supplementary trDTW experimentation***

I. *Average trDTW vs normalized DTW*

This simulation is to demonstrate that the average of the time-resolved DTW distances, obtained via PCHIP interpolation, closely approximates the normalized DTW distance. In each of the 10,000 realizations for the experiment, two independent white noise signals (i.i.d Gaussian, $\mathcal{N}(0,1)$) are generated, detrended, and then filtered using a low-pass Butterworth filter with a cutoff of 0.198 Hz. After z-score normalization, the DTW distance $D$ between the filtered signals is computed, along with the warping path indices, and then normalized by the length of the warping path.

The time-resolved DTW $D_{tr}\boldsymbol{(}\tau\boldsymbol{)}$ is computed using PCHIP interpolation to sample this back to the original signal length $D_{tr}^{pchip}\left( \tau\right)$. The average of $D_{tr}^{pchip}\left( \tau\right)$**,** $D_{tr-avg}$ is further computed.

The relative difference between the averaged $D_{tr-avg}$ and $D_{n}$ is then computed as

$$\left( \frac{\left| D_{tr-avg}-D_{n} \right|}{D_{n}} \right)\times100\%$$

The median relative difference over all realizations is found to be approximately 0.2%, indicating that the interpolated time-resolved DTW measure is nearly identical to the normalized DTW distance. This small discrepancy (0.2%) confirms that the PCHIP interpolation method provides an effective approximation of the normalized DTW in this context (Fig. 1).

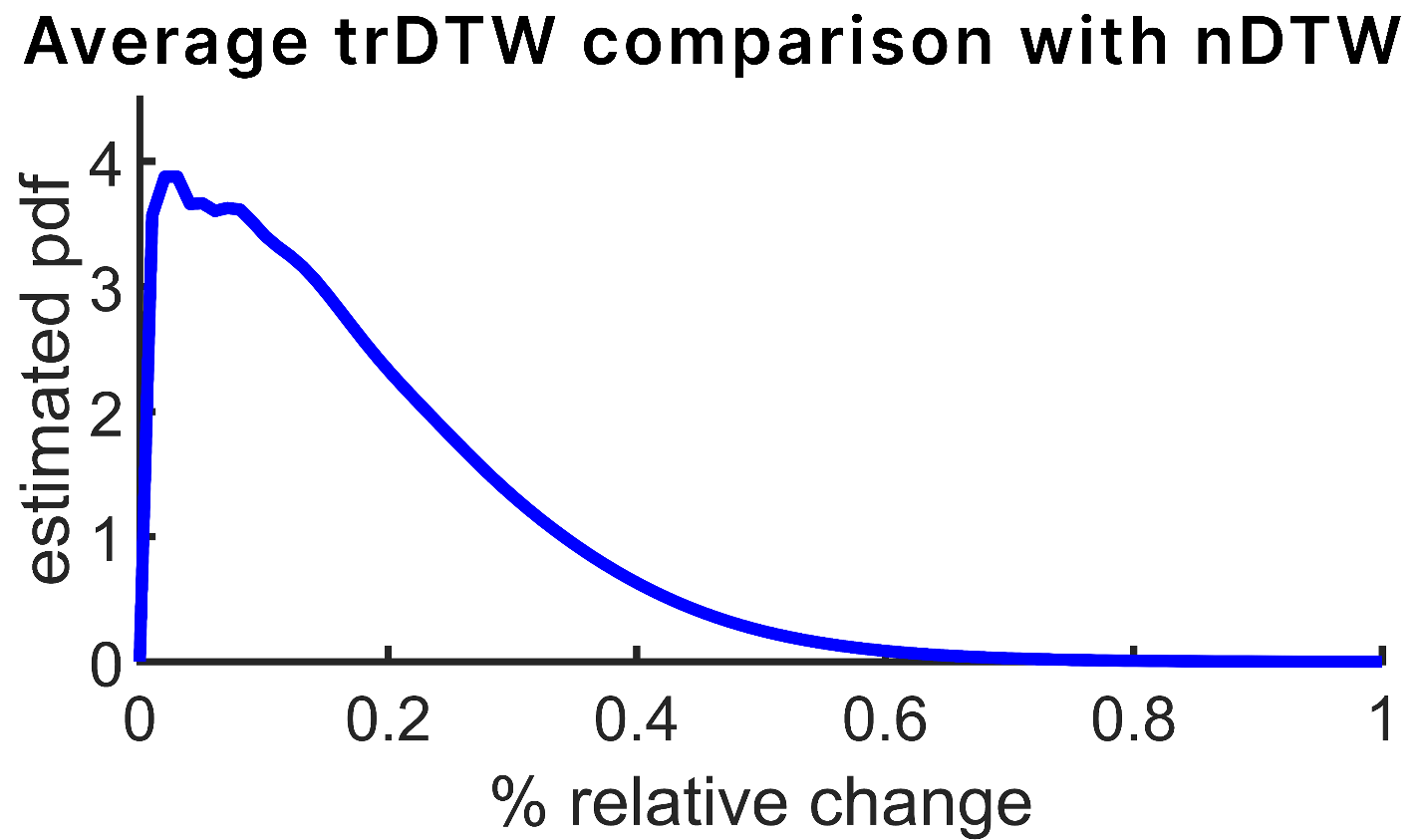

Figure 1. Comparison between the average time-resolved DTW and the normalized DTW.

***Supplementary brain states***

I. *Group differences in brain states*

After applying k-means clustering, the membership of each subject at each time point is extracted, allowing for the computation of subject representations within each cluster. A generalized linear model is then performed, controlling for age, sex, site, and mean frame displacement, to identify significantly different brain network pairs across the three brain states. To account for multiple comparisons, p-values are FDR-corrected within each brain state.

The results reveal only a few significant brain network pairs, particularly in the convergent and divergent states, when thresholded at 0.05 (Fig. 2). This suggests that these dynamic states are experienced in a highly similar manner across all subjects, including both controls and individuals with schizophrenia. As a result, the comparison of transitional dynamics across groups becomes more meaningful, as the underlying patterns are not entirely distinct.

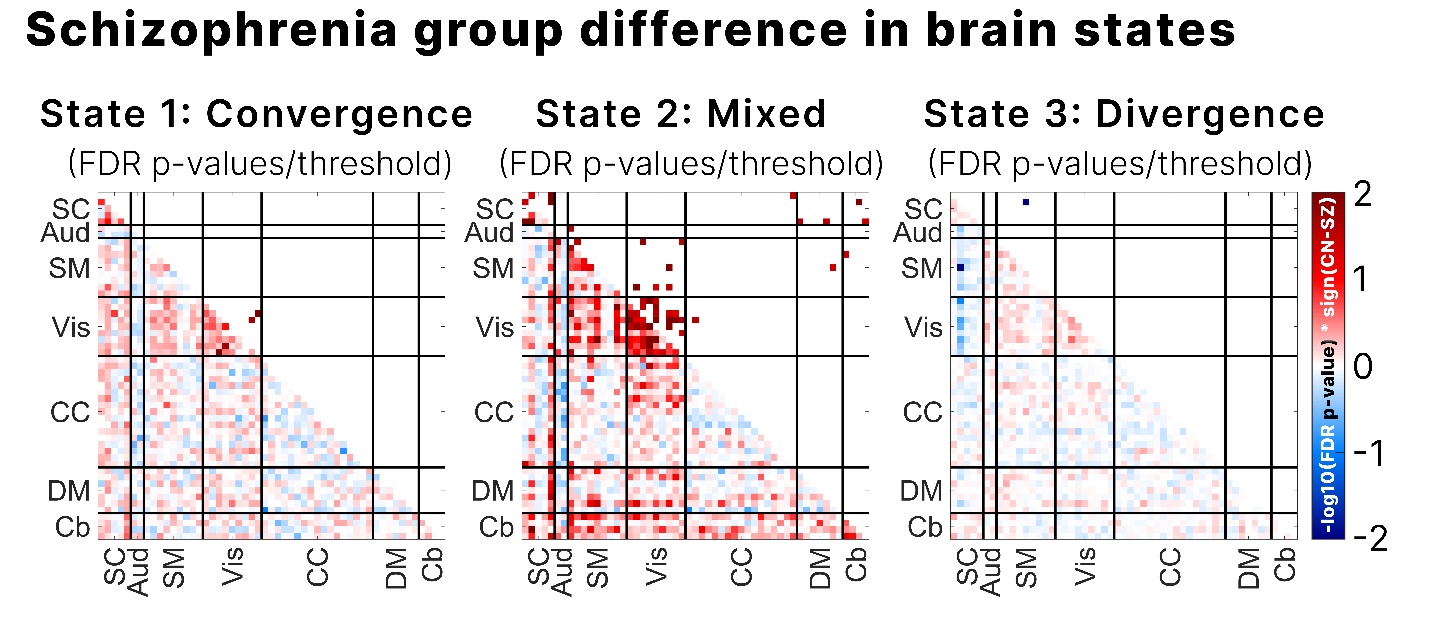

Figure 2 Group differences between schizophrenia and control groups for each brain state, derived from the time-resolved DTW analysis. A general linear model was used to assess group differences, controlling for sex, age, site, and mean frame displacement. FDR-corrected p-values are displayed in the lower triangle, while statistically significant results (FDR-corrected p < 0.05) are highlighted in the upper triangle.

***Supplementary directional DTW***

I. *Group differences in directional DTW*

Building on the derivation of the directional DTW measure, we computed the average directional DTW across groups to identify which brain networks exhibit greater amplitude relative to others. This analysis enabled us to assess the direction of disparities in metabolic energy demands across different brain regions.

Our findings indicate that while controls exhibit minimal amplitude disparity—suggesting a more convergent pattern—this disparity is primarily driven by a lower amplitude of the visual brain networks relative to the others. This suggests that, in controls, the visual brain region has a lower amplitude compared to other networks. The group differences presented in Fig. 3 further highlight that the observed disparity between the visual network and all other networks is largely due to the relatively lower amplitude of the visual network in controls, as opposed to schizophrenia patients.

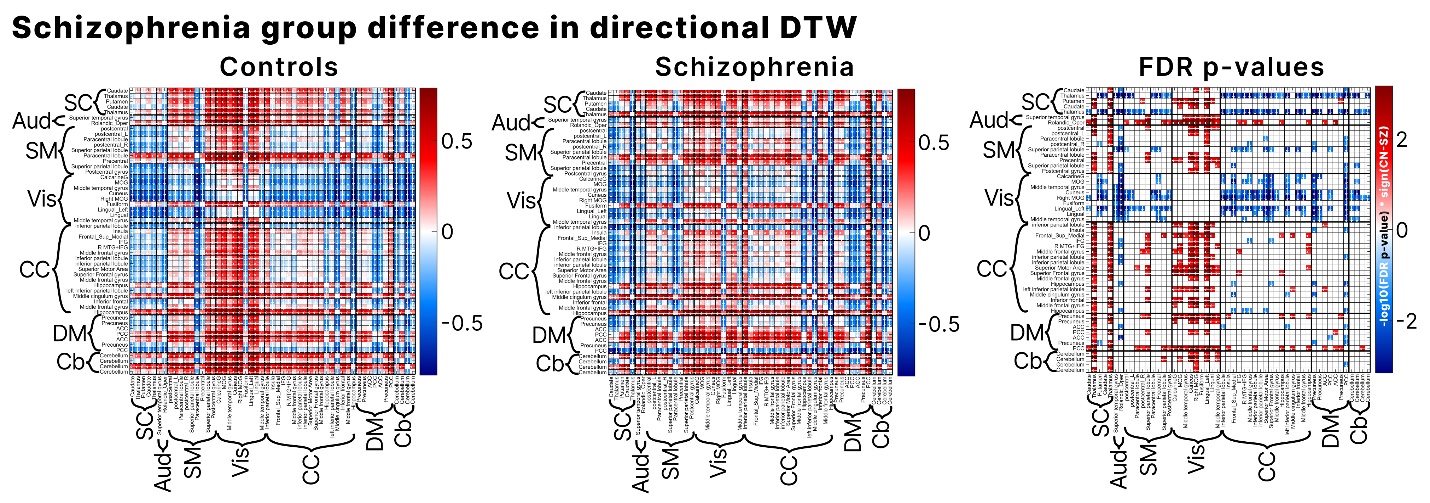

Figure 3 Group differences in the directional DTW metric between schizophrenia and control groups. The upper triangle is a sign-flipped version of the lower triangle for symmetry. Red cells indicate that the row network has a higher amplitude than the column network, while blue cells indicate that the column network has a higher amplitude than the row network. White arrows highlight the direction of the amplitude difference, with the arrow pointing toward the brain network that exhibits the highest amplitude.

II. *Brain state directional DTW*

We extracted directional time-resolved nDTW for each identified brain state, as depicted in Figure 4a. In these directional state matrices, red cells with white left-pointing arrows indicate that the brain network on the left has higher power than the one on the bottom, while blue cells with downward arrows signify that the brain networks at the bottom have higher power than those on the left. Our analysis reveals that the SC, Aud, and Cb domains consistently exhibit higher aligned power across all three states, whereas the Vis domain demonstrates lower aligned power relative to other networks.

Figure 4b illustrates the correlations between directional states, showing a high level of similarity. However, the distribution of power disparities between brain network pairs varies across states (Fig. 4c). A Kruskal-Wallis test confirmed that the distributions are significantly different (p < 0.05; Fig. 4d), indicating that while the directional information remains consistent, the magnitude of power differences determines whether a subject is in a convergent, divergent, or mixed state.

**Fig. 4: Directional time-resolved nDTW**

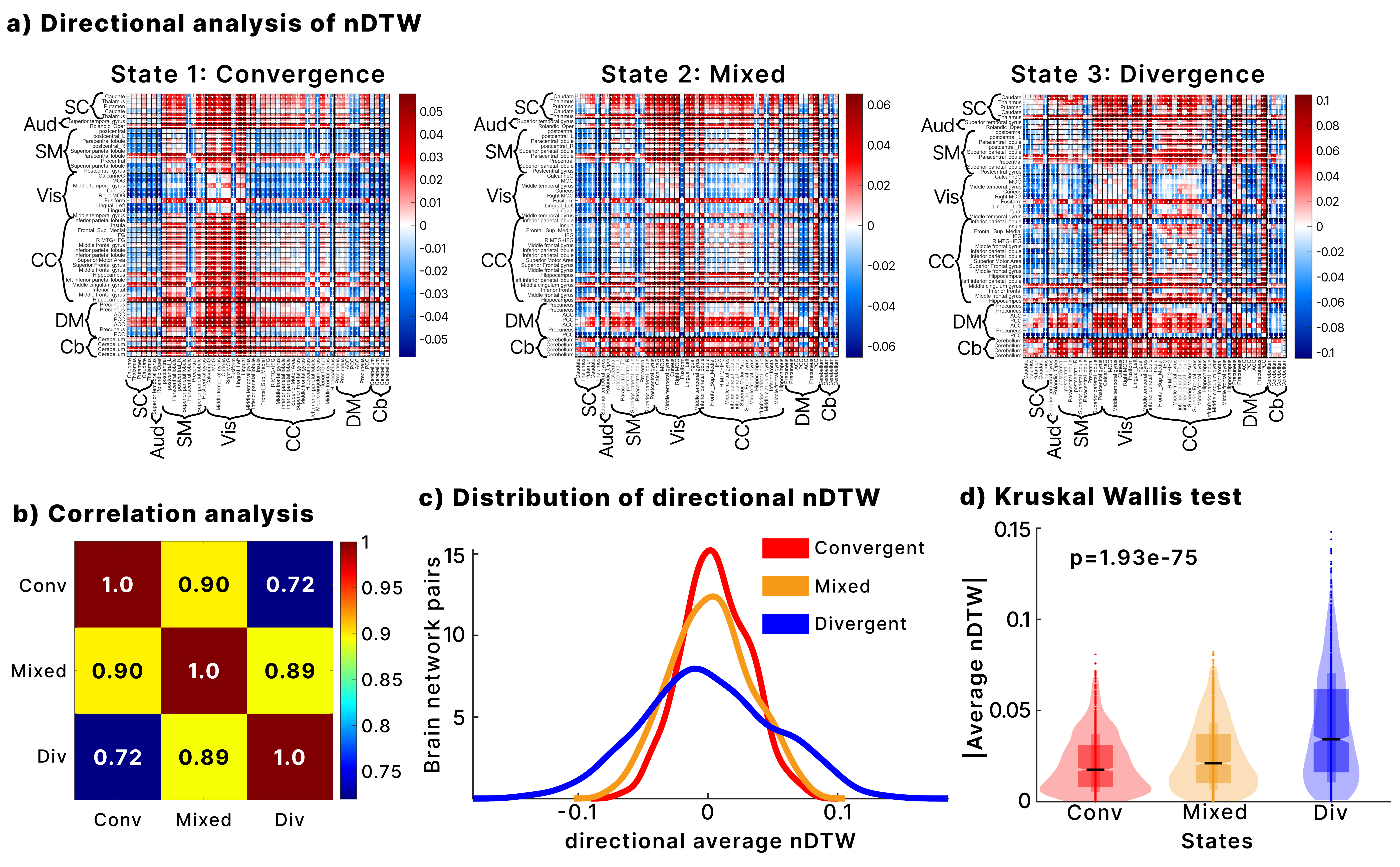
**a.** Directional nDTW matrices for each of the three identified brain states. Red cells with left-pointing arrows indicate higher power in left networks, while blue cells with downward-pointing arrows signify higher power in bottom brain networks. **b.** Correlation analysis between directional states, demonstrating similar patterns across states. **c.** Distribution of power disparities within each directional state, highlighting that the range of power imbalance varies across states. **d.** Kruskal-Wallis test results showing that the distributions of directional power disparities are significantly different among the three states (*p* < 0.05).

Our framework provides comprehensive insights into brain dynamics by analyzing BOLD signal changes—reflecting blood flow and oxygen levels as proxies for glucose utilization of the brain. Based on these insights, we propose that interventions for neurological disorders could aim to modulate whole-brain blood flow—using techniques such as photobiomodulation—to optimize transitions among convergent, divergent, and mixed states and thereby enhance cognitive performance. Notably, while the directional patterns of amplitude disparities across brain networks are consistent across all states (Fig. 4b), the magnitude of these disparities determines the current state. For example, to enhance attention vigilance in schizophrenia patients, controlled modulation targeted to enhance divergent to mixed state transitions should ensure that power disparities between brain networks move from a range of ~0.1 to a target range of ~0.06 (Fig. 4a). This targeted approach is intended to sustain the desired dynamic balance and promote optimal cognitive function.
